# Empagliflozin alleviates doxorubicin-induced cardiotoxicity by suppressing oxidative stress via modulation of the JNK/Nrf2 signaling pathway

**DOI:** 10.64898/2026.01.05.697595

**Authors:** Guangquan Peng, Lihua Ni, Ziyang Guo, Liwen Huang, Wanqing Ma, Fuchun Zheng, Yanmei Zhang, Fengfei Gao, Zhen Wang, Wenfeng Cai

## Abstract

Doxorubicin (DOX) treatment increases the risk of myocardial dysfunction and heart failure, in which oxidative stress plays a central role. Empagliflozin (EMPA) has been shown to benefit heart failure patients, yet its underlying mechanism remains unclear. In this study, we showed that EMPA treatment improved cardiac function, ameliorated structural remodeling, and increased survival in DOX-treated mice. Furthermore, EMPA reduced DOX-induced reactive oxygen species (ROS) overproduction and myocardial injury. Through functional analysis overlapping DIC- and EMPA-regulated genes, we identified the involvement of JNK signaling in addition to redox pathways. Subsequent investigations demonstrated that EMPA restored redox homeostasis by inhibiting JNK activation, reactivating NRF2 and its downstream antioxidant proteins HO-1, NQO1, GPX4, and SOD2 both *in vivo* and *in vitro*. These protective effects were abolished by the Nrf2 inhibitor ML385 or the JNK activator anisomycin. Collectively, our findings indicate that EMPA protects against DOX-induced cardiotoxicity by modulating the JNK/Nrf2 signaling and attenuating oxidative stress. This study provides a rationale for further investigation of EMPA as a potential therapeutic strategy for chemotherapy-associated cardiac injury.

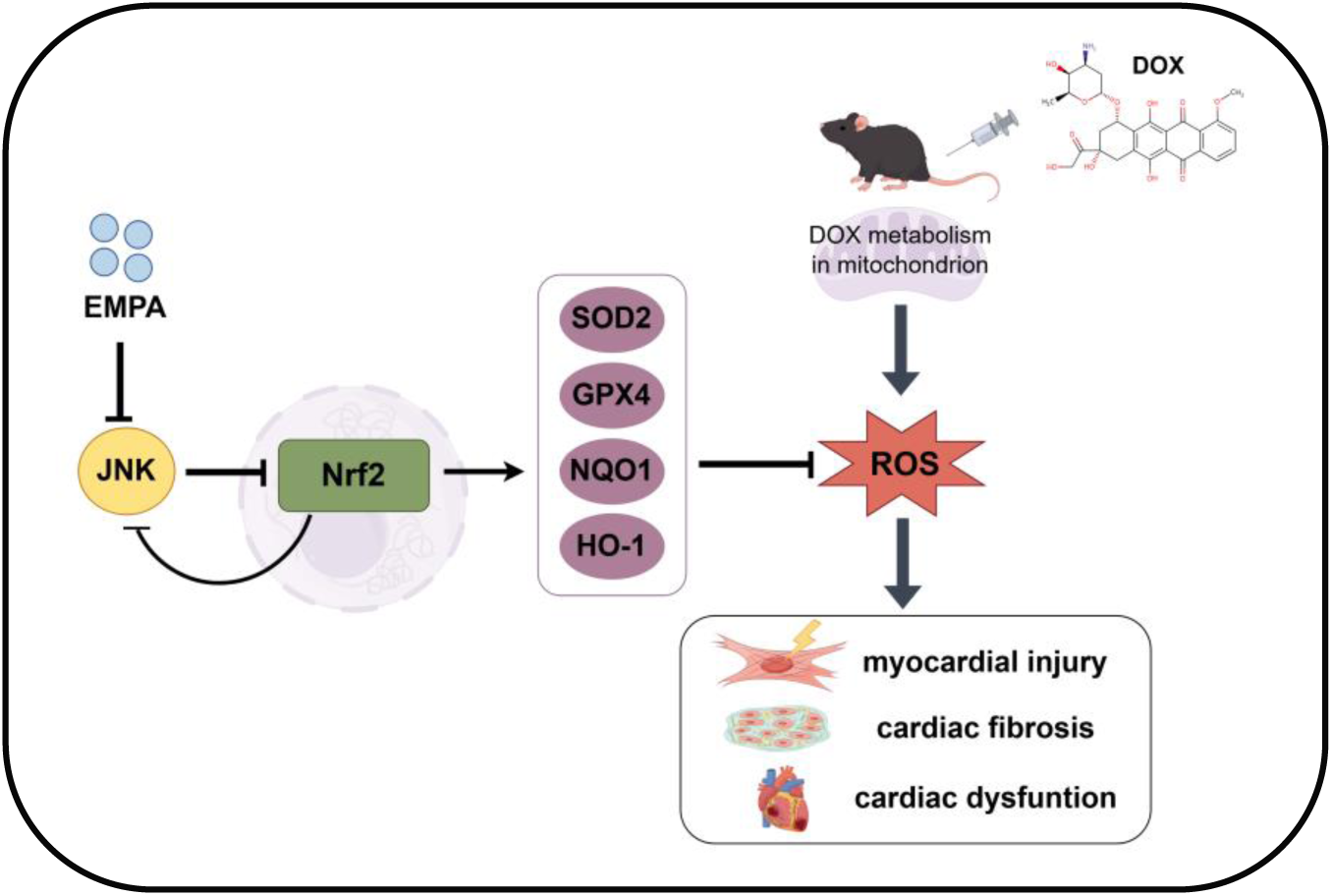

## 1. Introduction

Doxorubicin (DOX) is a highly effective anthracycline chemotherapeutic agent widely used in the treatment of various malignancies[1]. However, its clinical utility is significantly limited by dose-dependent cardiotoxicity, which can lead to irreversible myocardial dysfunction and heart failure[2]. The pathogenesis of DOX-induced cardiotoxicity (DIC) is multifactorial, with oxidative stress playing a central role. Upon entering cardiomyocytes, DOX undergoes enzymatic reduction in mitochondria to form semiquinone radicals, which react with molecular oxygen to generate excessive reactive oxygen species (ROS)[3,4]. This surge in ROS overwhelms cellular antioxidant defenses, resulting in lipid peroxidation, protein and DNA damage, mitochondrial dysfunction, and ultimately, cardiomyocyte death[5,6].

The cellular response to oxidative stress is tightly regulated by key signaling pathways. The transcription factor nuclear factor erythroid 2-related factor 2 (Nrf2) serves as a master regulator of antioxidant gene expression. Under physiological conditions, Nrf2 translocates to the nucleus and upregulates a network of cytoprotective enzymes, including heme oxygenase-1 (HO-1), NAD(P)H quinone dehydrogenase 1 (NQO1), glutathione peroxidase 4 (GPX4), and superoxide dismutase 2 (SOD2)[7,8]. In contrast, sustained oxidative stress can hyperactivate c-Jun N-terminal kinase (JNK), a member of the MAPK family, which promotes mitochondrial dysfunction, apoptosis, and further ROS accumulation[9,10]. Importantly, JNK activation has been shown to inhibit Nrf2 signaling, creating a vicious cycle that exacerbates oxidative injury[9,11]. Thus, the interplay between JNK and Nrf2 probably represents a critical regulatory node in DIC.

Empagliflozin (EMPA), a selective sodium-glucose cotransporter 2 (SGLT2) inhibitor, is well-established for its cardioprotective benefits in patients with type 2 diabetes and heart failure[12]. As increasing evidence indicates its cardiac protective effect can be independent of the glucose-lowering[13], European Heart Failure Guidelines recommend empagliflozin for HFrEF patients to improve symptoms, regardless of diabetes status[14]. A recent clinical study shows that SGLT2 inhibitors may also protect against chemotherapy-related cardiac dysfunction[15]. However, the precise molecular mechanisms through which EMPA mitigates DIC remain incompletely understood, particularly in non-diabetic contexts.

EMPA has been shown to activate antioxidant defenses[16]. It reduces ROS accumulation and suppress ventricular remodeling beyond metabolic regulation in both diabetic and non-diabetic models[12,17]. These indicate EMPA can modulate redox-sensitive pathways. Given this and the central role of oxidative stress in DIC, we hypothesized that EMPA protects against DOX-induced cardiotoxicity by restoring the balance between JNK and Nrf2 signaling. The purpose of this study is to 1) confirm EMPA alleviates DIC through inhibit oxidative stress; 2) investigate whether EMPA inhibit oxidative stress through interupting the JNK and Nrf2 feedback loop. Our findings will provide a novel mechanistic basis for EMPA cardioprotective action in the setting of chemotherapy-induced cardiac injury[18].

## 2. Materials and methods

### 2.1 Animal Procedure and Drug Treatment

All procedures were conducted according to the Institutional Animal Care Guidelines and were approved by the Animal Ethics Committee of the School of Medicine, Shantou University. Eight-week-old male C57BL/6J wild-type (WT) mice were randomly divided into two groups and received a single intraperitoneal injection of doxorubicin (DOX, 20 mg/kg) or an equivalent volume of normal saline (0.1 mL/10 g). Subsequently, the mice were orally administered empagliflozin (EMPA, 10 mg/kg/day) or vehicle (0.5% carboxymethyl cellulose sodium, 0.1 mL/10 g/day) once daily for seven consecutive days. At the end of treatment, all mice were euthanized, and heart tissues were collected for further analyses. Empagliflozin was kindly provided by Macklin (China) and prepared in 0.5% hydroxyethyl cellulose for oral gavage.

### 2.2 Echocardiographic Measurement

Cardiac function was evaluated using a Vevo LAZR high-resolution small-animal imaging system (VisualSonics, Toronto, Canada). Mice were anesthetized with isoflurane, and the inhalation rate was adjusted to maintain the heart rate between 400 and 500 beats per minute. Two-dimensional and M-mode echocardiographic recordings were obtained at the level of the left ventricular papillary muscles. Left ventricular ejection fraction (EF) and fractional shortening (FS) were calculated from M-mode tracings to assess systolic function.

### 2.3 Histological Examination

After seven days of treatment, mice were euthanized by intraperitoneal injection of an overdose of pentobarbital sodium. The hearts and tibias were collected for subsequent analyses. Cardiac tissues were fixed in 4% paraformaldehyde, dehydrated, embedded in paraffin, and sectioned at a thickness of 5–6 μm. Myocardial fibrosis was evaluated using Masson’s trichrome staining.

### 2.4 Dihydroethidium (DHE) Fluorescence Staining

Reactive oxygen species (ROS) levels in cardiac tissues were assessed by dihydroethidium (DHE) staining. Fresh-frozen heart sections (5 μm) were incubated with DHE solution at 37 °C for 30 min and then counterstained with DAPI using an antifade mounting medium. Fluorescence images were captured with a confocal laser scanning microscope (LSM 800).

### 2.5 Treatment of Cells and Experimental Design

H9c2 rat cardiomyocytes (ATCC, Rockville, MD, USA) were cultured in high-glucose DMEM supplemented with 10% FBS, 100 U/mL penicillin, and 100 μg/mL streptomycin at 37°C in a humidified 5% CO₂ atmosphere. Based on preliminary MTT assays, cells were exposed to doxorubicin (DOX, 1 μM) and empagliflozin (EMPA, 0.5 μM) for 24 hours according to the designated experimental groups, after which they were collected for downstream analyses.

### 2.6 Cell Viability Assay

Using the MTT assay following the manufacturer’s protocol. Briefly, H9c2 rat cardiomyocytes were seeded into 96-well plates at a density of 1 × 10⁴ cells per well and cultured under standard conditions. After the indicated treatments, 10 μL of MTT solution (5 mg/mL) was added to each well containing 100 μL of culture medium and incubated at 37 °C for 4 h. The supernatant was then carefully removed, and 150 μL of dimethyl sulfoxide (DMSO) was added to dissolve the formazan crystals. Absorbance was measured at 570 nm using a microplate reader.

### 2.7 Western Blotting

Heart tissues and H9C2 cells were lysed in RIPA buffer supplemented with protease and phosphatase inhibitors. Protein concentrations were quantified using a BCA protein assay kit. Equal amounts of protein were denatured in 5× loading buffer at 100 °C for 5 min, separated by 8–12% SDS–PAGE, and transferred onto nitrocellulose membranes. After blocking with 5% non-fat milk for 1 h at room temperature, the membranes were incubated with primary antibodies overnight at 4 °C followed by HRP-conjugated secondary antibodies for 2 h. GAPDH served as an internal control because of its stable expression and frequent use in cardiovascular research models. Protein bands were detected using an enhanced chemiluminescence imaging system (Bio-Rad) and quantified with ImageJ software.

### 2.8 Evaluation of Oxidative Stress Status

Oxidative stress markers, including malondialdehyde (MDA), superoxide dismutase (SOD), and glutathione (GSH/GSSG), were measured in serum and cell culture medium. Blood samples were collected from anesthetized mice via retro-orbital puncture, centrifuged at 3,000 × g for 10 min at 4°C, and serum was obtained. For cell experiments, culture medium was harvested for MDA and SOD analysis, while total GSH and GSSG were measured from cell lysates. MDA and SOD levels were determined using commercial ELISA kits (Nanjing Jiancheng Bioengineering Institute, Nanjing, China), with MDA quantified by the thiobarbituric acid (TBA) method (nmol/mL) and SOD activity assessed by the xanthine oxidase method (U/mL; one unit defined as the amount of enzyme required to inhibit 50% of the reaction rate). Total GSH and GSSG contents were analyzed using commercial assay kits (Beyotime Biotechnology, China) according to the manufacturer’s instructions.

### 2.9 LDH and CK-MB Assays

Lactate dehydrogenase (LDH) and creatine kinase-MB (CK-MB) activities were assessed in serum and H9c2 cardiomyocyte cultures. Blood samples were collected from anesthetized mice via retro-orbital puncture, centrifuged at 3,000 × g for 10 min at 4°C, and serum was obtained for analysis. Serum LDH and CK-MB activities were measured using a fully automated biochemical analyzer (Mindray BS-240, China) according to the manufacturer’s protocols. For cell experiments, LDH activity was determined in the culture medium using a commercial assay kit (Beyotime Biotechnology, China) following the manufacturer’s instructions. Results were expressed in U/L.

### 2.10 Intracellular ROS Detection

Intracellular reactive oxygen species (ROS) levels in H9c2 cardiomyocytes were assessed using the DCFH-DA fluorescent probe (Soleibao, China) according to the manufacturer’s instructions. Briefly, DCFH-DA was diluted 1:6000 in serum-free culture medium to prepare the working solution. After experimental treatments, cells were washed gently with PBS three times and incubated with 1 mL of the probe solution per well for 30 min at 37°C in the dark, with gentle shaking every 5 min. Cells were then washed three times with PBS and imaged under an inverted fluorescence microscope. The average fluorescence intensity of DCF was quantified using ImageJ.

### 2.11. Identification and functional enrichment of overlapping targets of DIC and EMPA

Chemical structure of EMPA were retrieved from PubChem and integrated from multiple pharmacological databases, including SwissTargetPrediction, CHEMBL, Super-Pred, PharmMapper, and the Comparative Toxicogenomics Database (CTD). Genes associated with DIC were collected from OMIM, GeneCards, and the Gene Expression Omnibus (GEO) dataset GSE157282. The overlap between the EMPA-associated target set and the DIC-related gene set was identified, yielding a list of candidate genes likely involved in EMPA’s cardioprotective effects.

Gene Ontology (GO) enrichment analysis was subsequently performed on these overlapping genes to characterize their functional profile. The analysis was conducted using the clusterProfiler R package (version 4.0). Significantly enriched terms were identified based on a hypergeometric test, with an adjusted p value < 0.05 and a q-value < 0.05 set as the significance thresholds.

### 2.12. Statistical analysis

Data are expressed as mean ± SEM. Statistical significance was defined as P < 0.05 (two-tailed). Comparisons between two groups were performed using Student’s t-test, while multiple group comparisons were analyzed by one-way ANOVA followed by Bonferroni’s post hoc test.

## 3. Results

### 3.1. Empagliflozin mitigates DOX-induced cardiac atrophy and dysfunction

To investigate the protective effects of empagliflozin (EMPA) against doxorubicin (DOX)-induced cardiotoxicity, wild-type (WT) mice were administered a single dose of DOX followed by daily oral treatment with EMPA for seven days. As shown in Figure 1A, DOX-treated mice exhibited marked cardiac atrophy, feature by significant reductions in the heart weight-to-tibia length (HW/TL) ratio (Figure 1A). EMPA treatment notably alleviated cardiac atrophy by improving HW/TL values. Kaplan–Meier survival analysis revealed a significantly higher survival rate in the EMPA-treated group compared with the DOX group (Figure 1B). Echocardiographic analysis revealed that DOX administration caused a pronounced decline in cardiac function, as indicated by reduced ejection fraction (EF) and fractional shortening (FS). EMPA treatment significantly restored both EF and FS toward normal levels (Figures 1C–E).

**Figure 1:**
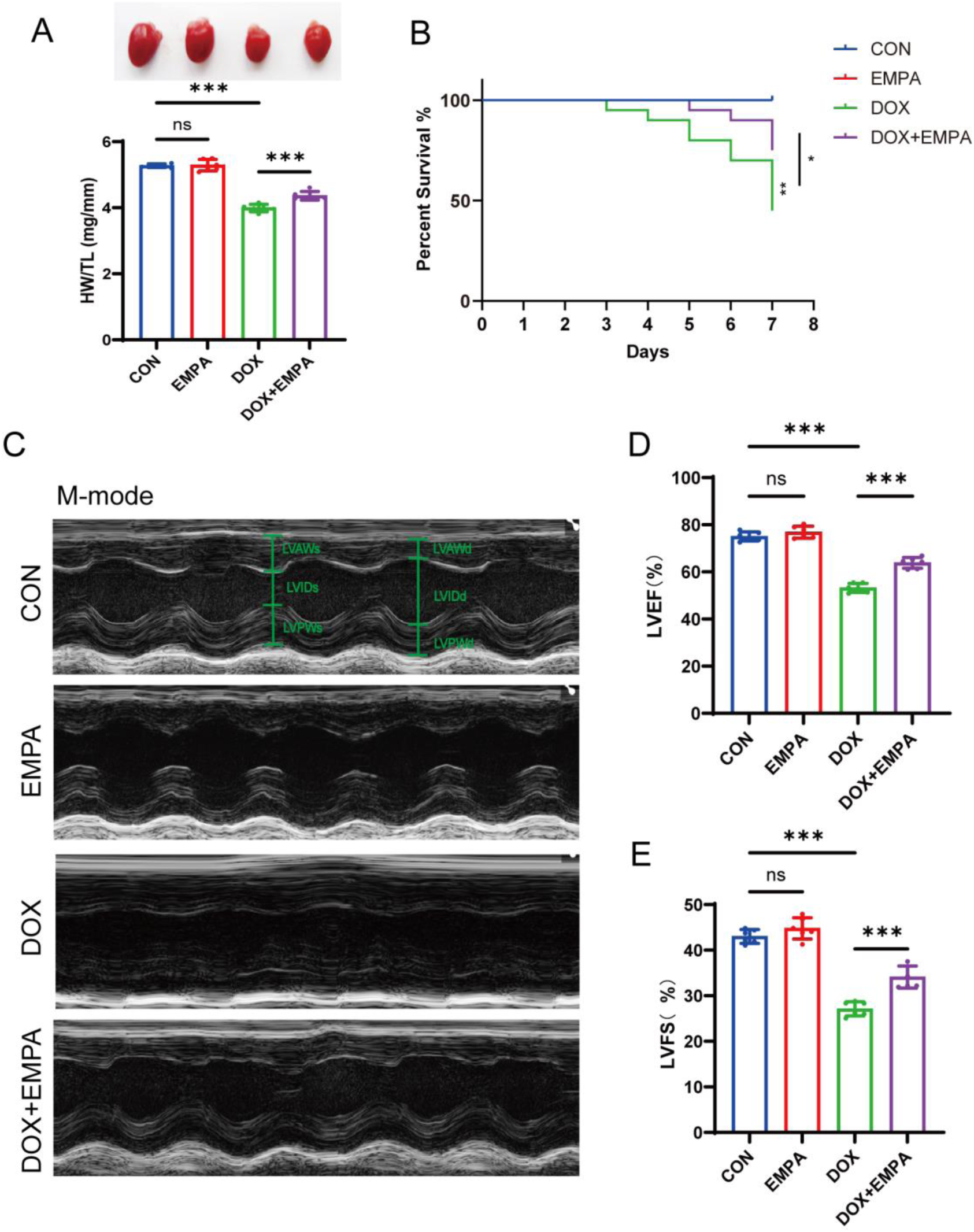
EMPA ameliorated DOX-induced cardiac atrophy and cardiac dysfunction in mice. (A) Representative images of mice hearts acorss groups and heart weight to body weight ratio (HW/BW), (B) Kaplan-Meier survival curves, (C) Representative echocardiographic M-mode images, (D) Left ventricular ejection fraction (EF), (E) Fractional shortening (FS). n = 6 per group for panels A, D, E, n = 20 per group for panel B. \**p* < 0.05, \*\**p* < 0.01, \*\*\**p* < 0.001.

### 3.2. EMPA alleviates DOX-induced myocardial fibrosis and injury

To further evaluate the histopathological alterations induced by DOX and the protective effects of EMPA, myocardial fibrosis was examined using Masson’s trichrome staining. DOX-treated mice exhibited extensive cardiac interstitial and perivascular collagen deposition, indicating significant myocardial fibrosis, whereas EMPA treatment markedly reduced collagen accumulation and preserved myocardial architecture (Figures 2A–B), Consistently, serum biochemical analyses revealed that DOX administration led to substantial increases in the cardiac injury markers lactate dehydrogenase (LDH), creatine kinase (CK), and its myocardial isoenzyme CK-MB. These elevations were significantly attenuated in the EMPA-treated group (Figures 2C–E), suggesting that EMPA effectively mitigates DOX-induced myocardial injury.

**Figure 2:**
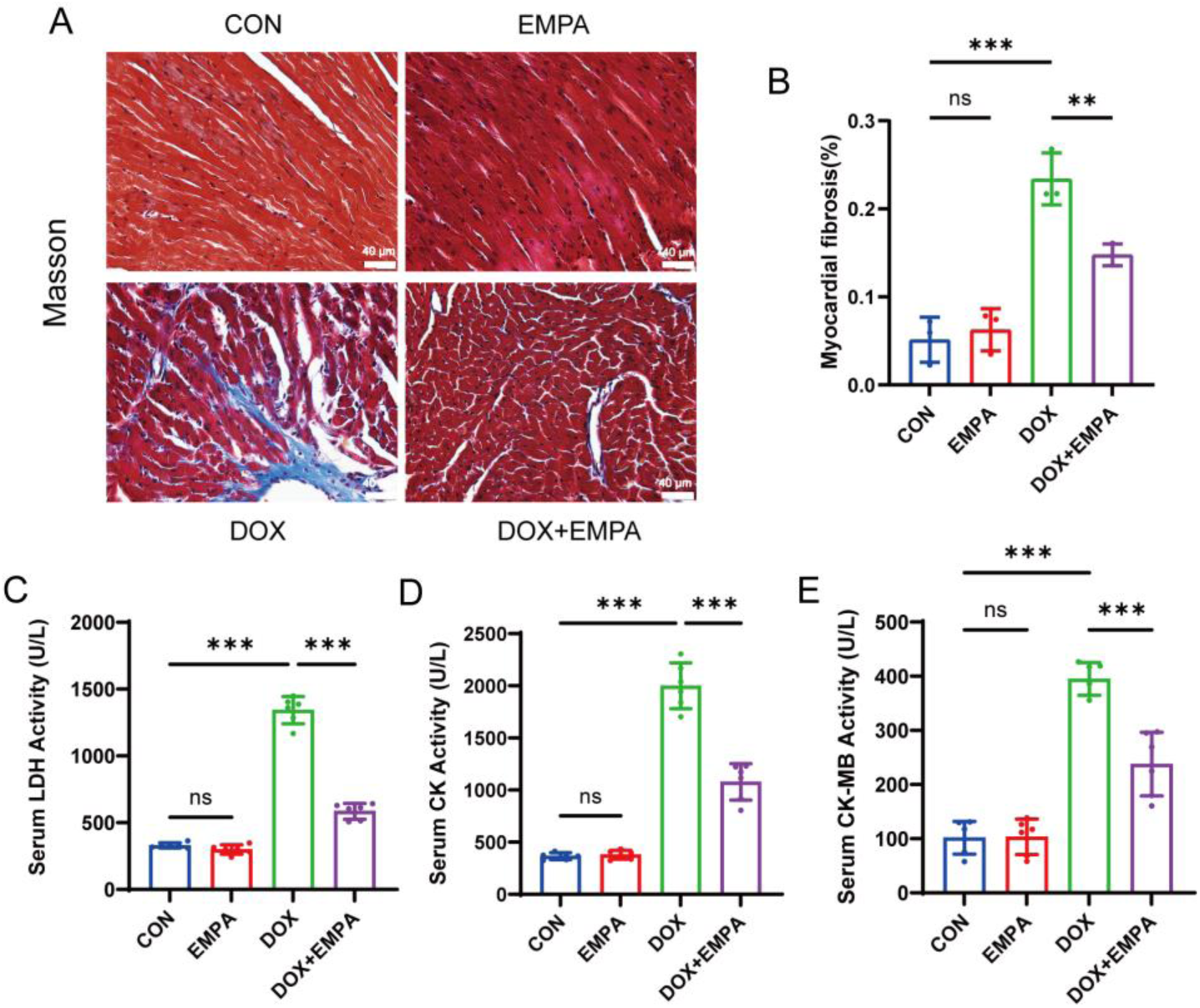
EMPA alleviated DOX-induced myocardial fibrosis and myocardial injury in mice. (A) Masson’s trichrome staining (collagen deposition in blue, scale bar = 40μm), (B) Quantiffcation of cardiomyocyte cross-sectional area, (C) Serum lactate dehydrogenase (LDH) activity, (D-E) Serum concentrations of creatine kinase (CK) and creatine kinase-MB isoenzyme (CK-MB). n = 3 per group for panels B, n = 6 per group for panel C-E. \**p* < 0.05, \*\**p* < 0.01, \*\*\**p* < 0.001.

### 3.3 EMPA suppresses oxidative stress in DOX-induced myocardium

To verify EMPA protects hearts from DIC through restore the redox balance, we next examined whether EMPA could attenuate DOX-induced oxidative stress in the myocardium. DHE staining was performed on frozen heart sections to detect reactive oxygen species (ROS) levels. DOX treatment markedly increased ROS accumulation in cardiac tissues, whereas EMPA administration significantly mitigated this increase (Figure 3A–B). Malondialdehyde (MDA), a lipid peroxidation product and a key marker of oxidative stress, was also substantially elevated in DOX-treated mice, which was significantly reduced by EMPA treatment (Figure 3C). Moreover, DOX administration decreased the glutathione (GSH) level and increased the oxidized glutathione (GSSG) level, resulting in a marked reduction in the GSH/GSSG ratio, indicating a disturbance of redox homeostasis. EMPA treatment reversed these alterations by increasing GSH and reducing GSSG levels, thus elevating the GSH/GSSG ratio (Figure 3D–F). Consistently, the activity of the antioxidant enzyme superoxide dismutase (SOD) suppressed by DOX exposure was restored by EMPA (Figure 3G). Collectively, these results demonstrate that EMPA effectively mitigates DOX-induced oxidative stress and restores antioxidant capacity in myocardium.

**Figure 3:**
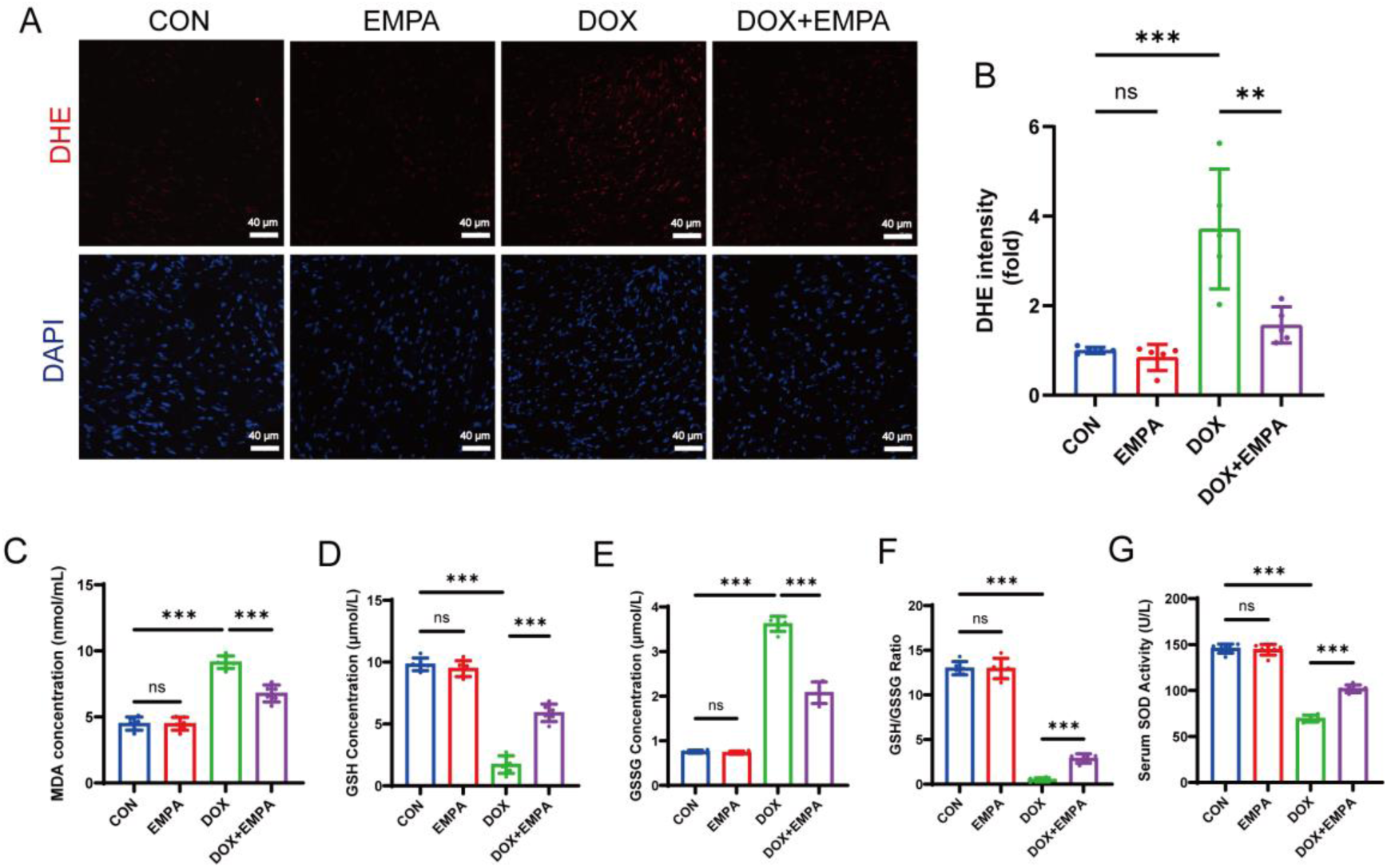
EMPA attenuated DOX-induced oxidative stress injury in mouse myocardium. (A) Representative images of DHE staining in cardiac sections (red, ROS; blue, nuclei; scale bar = 40 μm), (B) quantitative analysis of ROS in cardiac tissues, (C) serum MDA level, (D–F) serum GSH concentration, GSSG concentration, and the GSH/GSSG ratio, (G) serum SOD level. n = 5 per group for panels B, n = 6 per group for panel C-G. \**p* < 0.05, \*\**p* < 0.01, \*\*\**p* < 0.001.

### 3.4 EMPA attenuates DOX-induced oxidative stress in H9c2 cells

To further confirm the in vivo findings and directly assess cardiomyocyte responses, we evaluated the effects of EMPA in H9c2 cells. Consistent with the animal data, DOX exposure significantly elevated intracellular ROS levels and MDA content, while depressing SOD activity and disrupting the GSH/GSSG ratio (Figure 4A–G). EMPA co-treatment effectively reversed all these oxidative stress parameters. Moreover, EMPA markedly reduced DOX-induced lactate dehydrogenase (LDH) release, a direct indicator of cardiomyocyte membrane integrity and viability (Figure 4H). These results confirm that EMPA exerts direct antioxidative and cytoprotective effects against DOX insult in cardiomyocytes.

**Figure 4:**
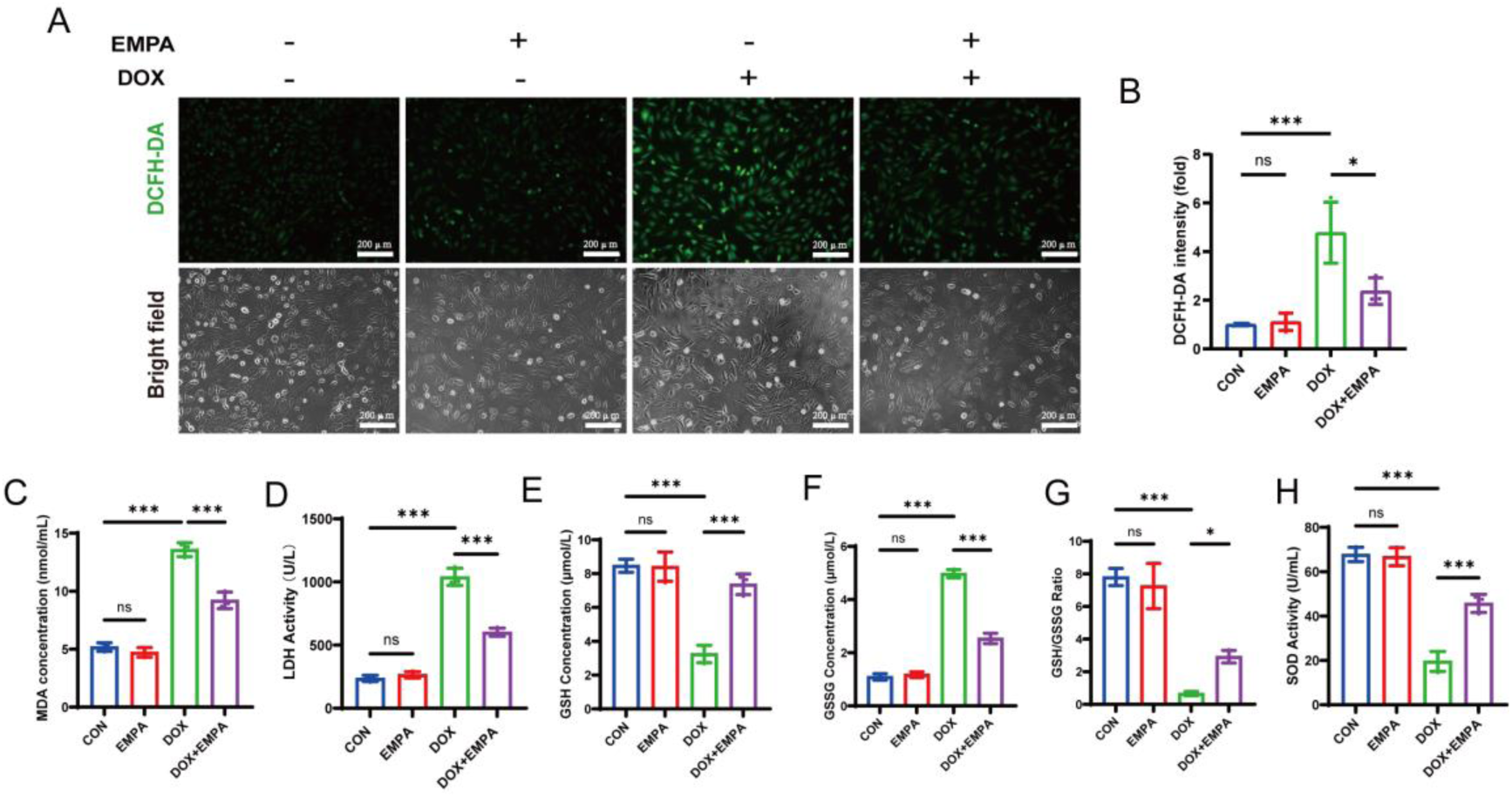
EMPA attenuated DOX-induced oxidative stress in H9c2 cells. (A) Representative graph of DCFH-DA staining of H9c2 cells, (B) Quantitative analysis of ROS content, (C) MDA content in the culture medium of H9c2 cells, (D) The LDH activity, (E-G) The concentrations of GSH and GSSG in H9c2 cell lysates, as well as the GSH/GSSG ratio, (H) SOD content in the culture medium of H9c2 cells. n = 3 per group. \**p* < 0.05, \*\**p* < 0.01, \*\*\**p* < 0.001.

### 3.5 Transcriptomic Analysis Implicates Oxidative Stress and JNK/NRF2 Pathways

The result above demonstrated that EMPA effectively alleviated DOX-induced myocardial injury. To further explore the underlying mechanisms of DOX-induced cardiotoxicity and the protective effects of EMPA, we analyzed the overlap targets of DIC and EMPA. Totally, we identified 236 targets associated with EMPA and 2682 DIC-related genes from public database. Among them, 140 overlapping genes were identified. The oxidative stress related signaling pathway were enriched in both biological process (BP) and molecular function (MF) items. Notably, the JNK pathway were highlighted in both BP and MF section.

**Figure 5:**
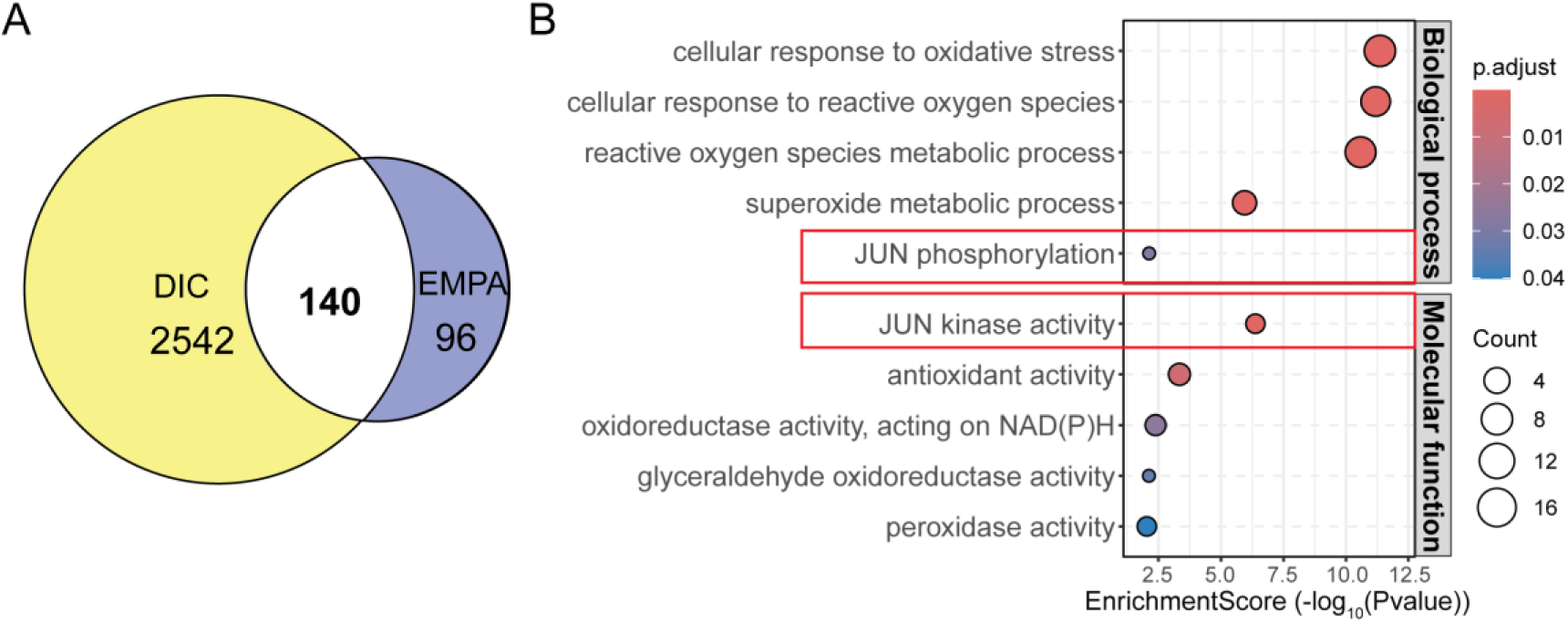
A. The overlap of DIC regulated genes and EMPA associated targets identified. B. GO analysis for the 140 overlapped targets. Redbox indicates the JNK pathway.

### 3.6. EMPA Modulates the JNK/Nrf2 Signaling Pathway in vivo

Given that the transcriptomic analysis highlighted the JNK and NRF pathways, we next focused on its potential involvement in EMPA-mediated protection against DOX-induced oxidative stress. We examined the protein expression levels of these signaling molecules in myocardial tissues by Western blot analysis. DOX treatment markedly increased the phosphorylation of JNK, and decreased the expression of Nrf2, HO-1, and their downstream antioxidant proteins NQO1, SOD2, and GPX4. In contrast, EMPA administration significantly attenuated JNK hyperphosphorylation and restored the expression of Nrf2 and its downstream antioxidant effectors (Figure 6A–G). These findings support our hypothesis that EMPA mitigates DOX-induced oxidative stress injury, at least in part, by suppressing JNK overactivation and enhancing the Nrf2/HO-1 antioxidant signaling pathway.

**Figure 6.**
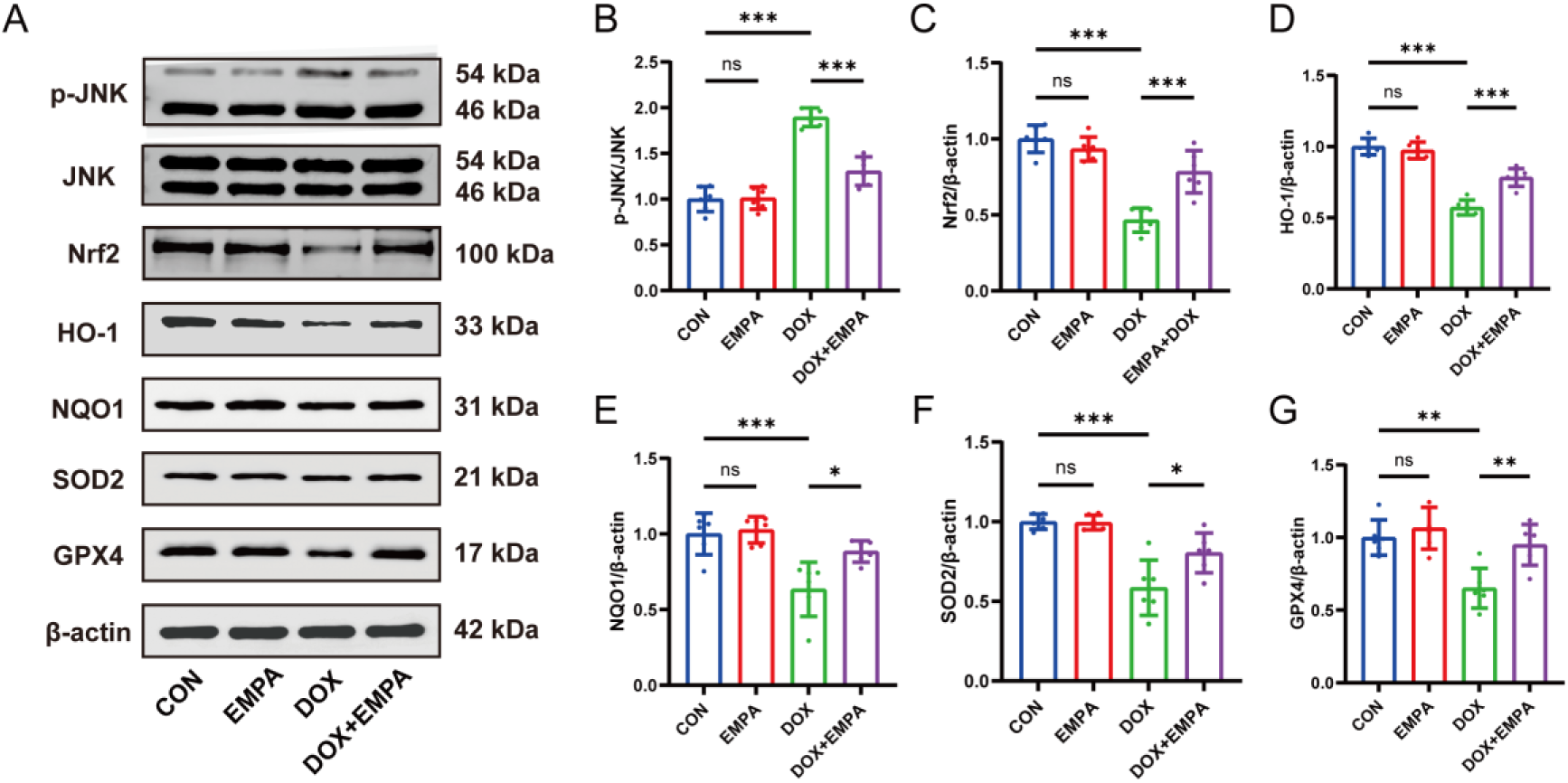
EMPA modulated the JNK and Nrf2/HO-1 signaling pathways in DOX-induced myocardial tissues. (A–G) Western blot images and semi-quantitative analyses of JNK, Nrf2, HO-1, NQO1, SOD2, and GPX4 proteins. n = 6 per group. \**p* < 0.05, \*\**p* < 0.01, \*\*\**p* < 0.001.

### 3.7 EMPA Modulates the JNK/Nrf2 Signaling Pathway in vitro

We next confirmed these signaling effects in H9c2 cardiomyocytes. Consistent with the in vivo results, DOX exposure induced pronounced JNK phosphorylation and downregulated Nrf2, HO-1, NQO1, SOD2, and GPX4 protein levels. EMPA pre-treatment effectively inhibited JNK activation and restored the expression of Nrf2 and its downstream effectors (Figures 7A-G). These in vitro data reinforce the conclusion that EMPA targets the JNK/Nrf2 axis to counteract DOX-induced oxidative stress.

**Figure 7:**
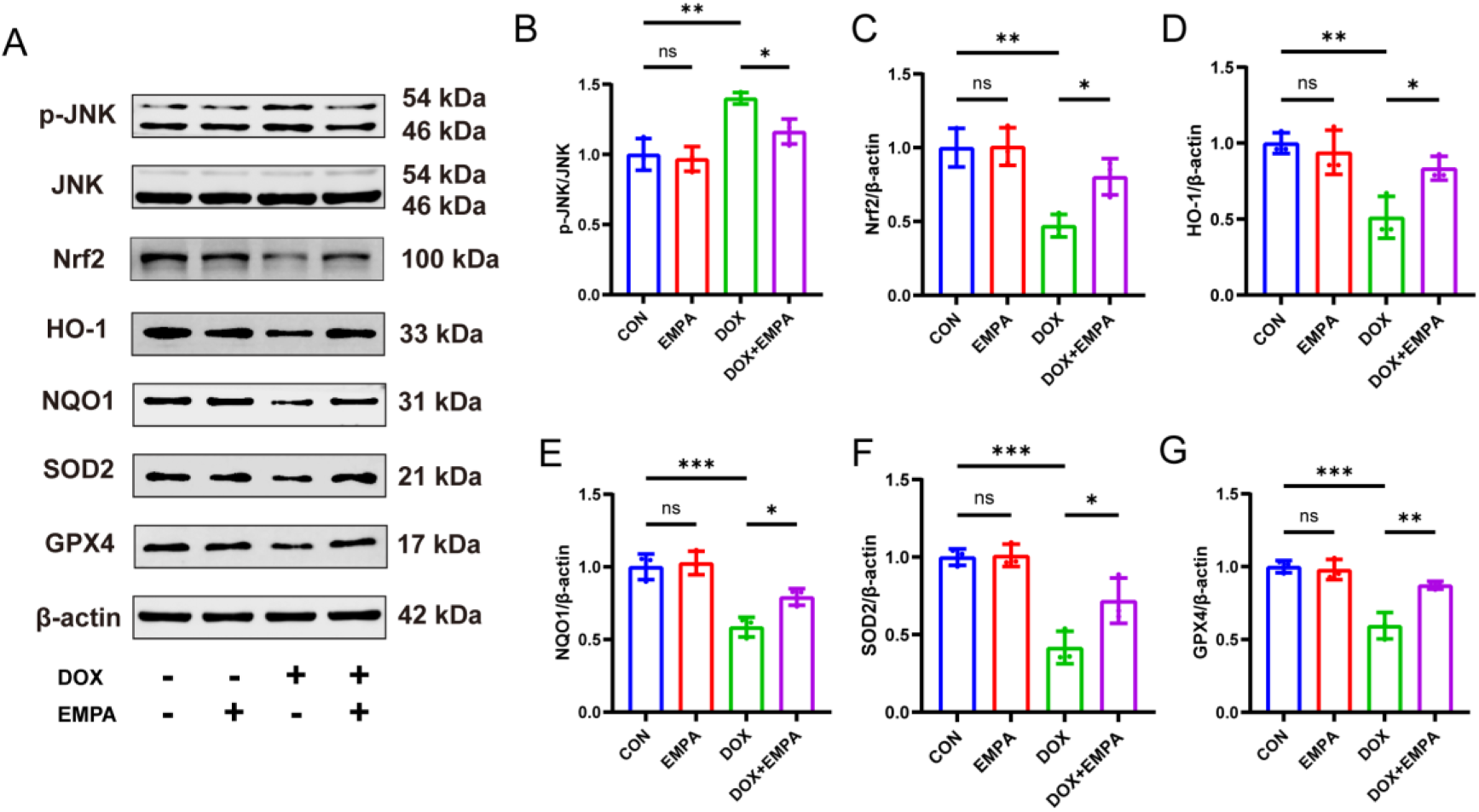
EMPA modulated the JNK and Nrf2/HO-1 signaling pathways in DOX-induced H9C2 cardiomyocytes. (A–G) Western blot images and semi-quantitative analyses of JNK, Nrf2, HO-1, NQO1, SOD2, and GPX4 proteins. n = 3 per group. \**p* < 0.05, \*\**p* < 0.01, \*\*\**p* < 0.001.

### 3.8. JNK re-activation abolished the protective effects of EMPA

To further confirm that JNK is involved in EMPA-mediated antioxidant effects and its regulation of Nrf2, we performed intervention experiments at the cellular level using the JNK activator Anisomycin. Anisomycin treatment reversed the suppressive effect of EMPA on DOX-induced ROS accumulation, leading to significantly higher ROS levels compared to the DOX+EMPA group (Figure 8A–B). Western blot analysis further revealed that Anisomycin enhanced JNK phosphorylation while significantly reducing Nrf2 expression in the presence of EMPA (Figure 8C-G). These findings indicate that EMPA may exert its antioxidant effects by suppressing JNK hyperactivation and suggest a negative regulatory role of JNK on Nrf2.

**Figure 8:**
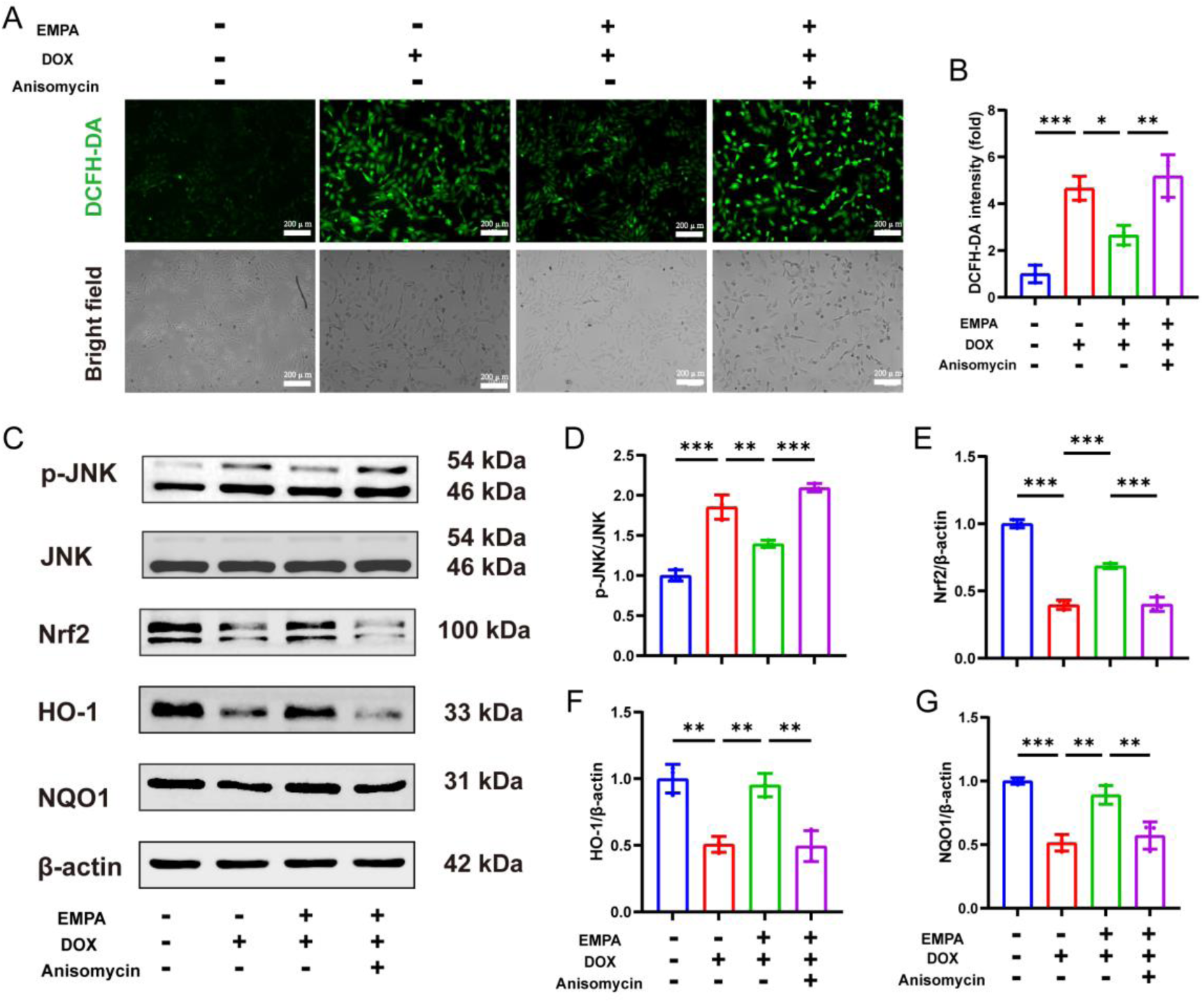
Role of JNK in EMPA-mediated regulation of myocardial resistance to oxidative stress. (A) Representative graph of DCFH-DA staining in H9c2 cells; (B) Quantitative analysis of ROS content; (C-G) Western Blot for protein immunoblotting and semi-quantitative analysis of JNK, Nrf2, HO-1, and NQO1. n = 3 per group. \**p* < 0.05, \*\**p* < 0.01, \*\*\**p* < 0.001.

### 3.9. Inhibition of Nrf2 abolishes EMPA-mediated antioxidative protection

To confirm Nrf2 activation is essential in mediating EMPA’s antioxidative effect, we employed the specific Nrf2 inhibitor ML385. ML385 treatment effectively abolished EMPA’s suppression of DOX-induced ROS accumulation (Figures 9A, B). Furthermore, ML385 not only decreased the expression of Nrf2 and its target proteins (HO-1, NQO1) but also concurrently enhanced JNK phosphorylation, even in the presence of EMPA (Figures 9C-G). These findings demonstrate that Nrf2 activation I s required for EMPA’s cytoprotection and reveal a reciprocal negative regulation between Nrf2 and JNK signaling in this context.

**Figure 9:**
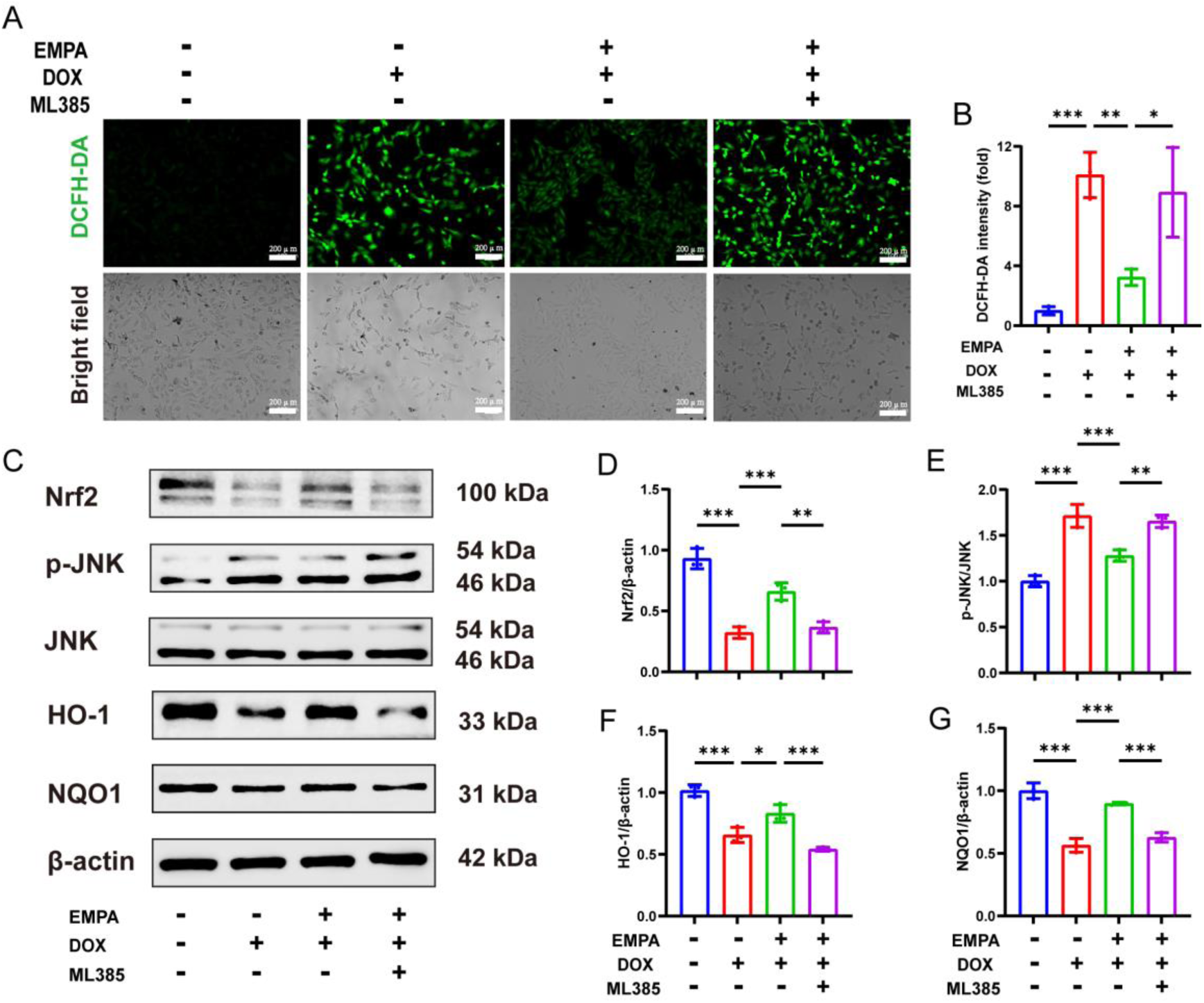
Role of Nrf2 in EMPA-mediated regulation of myocardial resistance to oxidative stress. (A) Representative graph of DCFH-DA staining in H9c2 cells, (B) Quantitative analysis of ROS content, (C-G) Western Blot for protein immunoblotting and semi-quantitative analysis of JNK, Nrf2, HO-1, and NQO1. n = 3 per group. \**p* < 0.05, \*\**p* < 0.01, \*\*\**p* < 0.001.

## Discussion

In this study, we demonstrated that empagliflozin confers robust protection against dox-induced cardiotoxicity using both *in vivo* and *in vitro* model. We showed that EMPA attenuates DOX-induced cardiac atrophy, preserves systolic function, reduces circulating markers of myocardial injury, and markedly suppresses oxidative stress. Mechanistically, EMPA inhibited JNK hyperphosphorylation triggered by DOX and restored Nrf2-mediated antioxidant responses (HO-1, NQO1, SOD2, GPX4). Notably, the cardiac protective effect of EMPA was abolished by reactivation of JNK or inhibition of Nrf2, further validating that EMPA mitigates DOX-induced oxidative stress and cardiac injury by targeting the JNK-Nrf2 feedback loop

DOX-induced cardiac dysfunction is characterized by protein/DNA damage and cell death at the cellular level, and clinically by myocardial atrophy and progressive ventricular dysfunction[19,20]. Our model, employing a single high-dose DOX injection, successfully recapitulated these key features—evidenced by a reduced heart weight-to-tibia length ratio, elevated serum levels of LDH, CK, and CK-MB, and depressed LVEF and LVFS—confirming that the experimental phenotype reliably mirrors DOX-induced injury.

Oxidative stress and mitochondrial dysfunction are central initiating events in DIC, leading to lipid peroxidation and cellular dysfunction[21]. Our results consistently show that DOX administration induces a profound redox imbalance, characterized by excessive ROS generation, elevated MDA, depleted GSH, and suppressed SOD activity in both myocardial tissue and H9c2 cardiomyocytes.

Currently, effective strategies to prevent or treat DOX-induced cardiotoxicity are limited. Since 2021, EMPA has been recommended in guidelines to improve outcomes in patients with heart failure with reduced ejection fraction (HFrEF), irrespective of diabetes status[22]. In 2022, Gongora et al. provided the first clinical evidence that SGLT2 inhibitor use is associated with a lower risk of cardiac dysfunction in patients receiving anthracyclines, although the underlying mechanism remained unidentified[15]. Our study confirms the direct cardioprotective effect of EMPA against DIC in a preclinical model, supporting these clinical observations and strengthening the rationale for further investigation of EMPA’s role in cardio-oncology.

EMPA has been shown to reduce ROS accumulation, attenuate inflammation, and suppress adverse remodeling in various cardiac injury models, both diabetic and non-diabetic[17,23,24]. However, its specific effect on DOX-induced oxidative stress was unclear. We found that, consistent with its role in other settings, EMPA potently inhibits DOX-induced oxidative stress. Treatment effectively reversed all measured oxidative stress markers in DIC, restoring the GSH/GSSG ratio and SOD activity. Furthermore, EMPA actively enhanced antioxidant defenses, aligning with observations of its antioxidant properties in models of aging[25,26]. Together, this demonstrates that EMPA possesses significant antioxidant capacity, which likely forms a cornerstone of its cytoprotective action against DOX.

At the molecular level, we delineated a critical signaling interplay regulated by EMPA to against DIC. We showed that DOX treatment induced sustained phosphorylation/activation of JNK, a stress-activated kinase known to promote apoptosis and mitochondrial dysfunction, while simultaneously suppressing the Nrf2 pathway and its downstream antioxidants (HO-1, NQO1, SOD2, GPX4). EMPA administration rectified this imbalance by inhibiting JNK activation and promoting Nrf2 target gene expression. Notably, JNK activator anisomycin and the Nrf2 inhibitor ML385 abolished the cardiac protective effect of EMPA, confirmed the important roles of these pathways. Interestingly, ML385 not only blocked Nrf2 activation but also reversed EMPA’s suppression of JNK, suggesting a negative feedback loop where Nrf2 activation helps restrain JNK signaling. This is consistent with previous findings that NRF2’s activation negative regulated JNK hyperactivation[11,27].

## Conclusions

our study demonstrates that empagliflozin provides significant protection against acute doxorubicin-induced cardiotoxicity by mitigating oxidative stress through modulation of the JNK-Nrf2 interplay pathways. These findings elucidate a novel molecular mechanism underlying EMPA’s cardioprotective action and provide strong preclinical justification for its potential application in cardio-oncology. By targeting a central pathological process in DIC with a drug already proven safe for long-term clinical use, this research highlights a promising therapeutic strategy to enhance the safety and sustainability of anthracycline-based chemotherapy.

## Limitations

Our study has several limitations. First, the protective effects of EMPA were evaluated in an acute model; its efficacy in a chronic, low-dose DOX exposure model mimicking clinical chemotherapy cycles warrants further investigation. Second, while we focused on the JNK/Nrf2 axis and oxidative stress, EMPA may also exert benefits through other pathways, such as improving mitochondrial energetics, modulating inflammation, or inhibiting ferroptosis, which were not explored here.

## CRediT authorship contribution statement

W.C. and Z.W. conceived the project. G.P and L.N. performed the experiments, analyzed, and compiled the data. Z.G., L.H., W.M., F.Z., Y.Z., and F.G. helped with experiments. G.P., Z.W., and W.C. participated in writing or editing the paper.

## Data availability

The data that support the findings of this study are available on request from the corresponding author.

